# Uncovering plant microbiomes using long-read metagenomic sequencing

**DOI:** 10.1101/2023.03.20.533568

**Authors:** Sachiko Masuda, Pamela Gan, Yuya Kiguchi, Mizue Anda, Kazuhiro Sasaki, Arisa Shibata, Wataru Iwasaki, Wataru Suda, Ken Shirasu

## Abstract

The microbiome of plants plays a pivotal role in their growth and health. Despite its importance, many fundamental questions about the microbiome remain largely unanswered, such as the identification of colonizing bacterial species, the genes they carry, and the location of these genes on chromosomes or plasmids. To gain insights into the genetic makeup of the rice leaf microbiome, we performed a metagenomic analysis using long-read sequences, and developed a genomic DNA extraction method that provides relatively intact DNA for long-read sequencing. 1.8 Gb reads were assembled into 26,067 contigs, including 136 circular sequences of less than 1 Mbp, as well as 172 large (≥ 1 Mbp) sequences, six of which were circularized. Within these contigs, 669 complete 16S rRNA genes were clustered into 166 bacterial species, 130 of which showed low identity to previously defined sequences, suggesting that they represent novel species. The large circular contigs contain novel chromosomes and a megaplasmid, and most of the smaller circular contigs (<1 Mbp) were defined as novel plasmids or bacteriophages. One circular contig represents the complete chromosome of an uncultivated bacterium in the candidate phylum *Candidatus Saccharibacteria*. Our findings demonstrate the efficacy of long-read-based metagenomics for profiling microbial communities and discovering novel sequences in plant-microbiome studies.

## Introduction

Plants provide a wide range of ecological niches for microbes, including the surfaces of organs such as leaves, flowers, fruits and roots, internal structures that are colonized by endosymbionts, the zone of intimate interactions between plant roots and microbes known as the rhizosphere, and into the soil beyond the rhizosphere due to the diffusion of plant products away from the root zone. These diverse environments allow microbes to form complex communities collectively known as the plant microbiome^1^ or phytobiome^2^ that can promote plant growth by affecting nutrient uptake, by suppressing pathogens, and by inducing disease resistance^3^. While individual members of the plant microbiome may express beneficial traits, because it is a complex system, the overall effects of the microbiome on plant health cannot be predicted from individual microbial taxa^3^. The community composition of a plant microbiome is influenced by host plant genetics^4, 5^, specificity and activity of plant defenses^6^, microbe-microbe interactions^7^, and environmental factors such as soil geochemistry and UV light intensity. Understanding the mechanisms by which the plant microbiome impacts plant health is a crucial area of research, with the potential to inform the development of new strategies for improving plant growth, productivity, and sustainability.

Plasmids and bacteriophages and the mobile genetic elements they carry, including IS elements and transposons, have a role in shaping microbial communities by providing novel capabilities^8, 9^, such as the well-studied VirB/VirD4 system found in both pathogenic and symbiotic bactera in plant-microbiome interactions^10–13^. The VirB/VirD4 system, typcially encoded on a plasmid, enables pathogenic bacteria to invade host plants by transporting effector proteins and manipulating the host’s immune system. On the other hand, the VirB/VirD4 system in root-nodulating bacteria is involved in protein translocation and can have a host-dependent effect on symbiosis^14^. Despite their ecological importance, our understanding of plasmid and bacteriophage sequences from isolated bacteria is limited.

To gain deeper insights into the ecological and biological features of plant microbiomes, culture-independent metagenomics has increasingly been employed as a tool to analyze the genetic makeup of complex microbial communities^1^. This approach provides a catalog of the microbial diversity and functional potential within a given community^1^. However, traditional short-read (<500 bp) sequencing, typically of one or two of the variable regions of 16S rRNA genes or other single genes, poses serious limitations for accurate identification and genome reconstruction^15, 16^. Primer sequence bias, and a large proportion of short reads that cannot be mapped to a reference genome results in the loss of potentially useful information^17^. In contrast, long-read metagenomics has the potential to generate longer contigs, thus improving genome reconstruction, taxonomic assignment, and revealing previously undiscovered sequences, including circular genomes and extrachromosomal elements. For instance, long-read metagenomic sequencing of the human gut microbiome has uncovered a higher number of plasmids than previously reported^18^. We anticipate that long-read metagenomics will be a valuable approach for exploring plant microbiome metagenomes.

In this study, we used long-read metagenomics to better understand the genetic makeup of the rice (*Oryza sativa*) microbiome. We enriched microbes from the phyllosphere and established a genomic DNA extraction method for long-read sequencing. Then, using reads from the Pacbio Sequel II sequencer, we reconstructed 26,067 contigs, including novel circularized chromosomes, plasmids and bacteriophages. Notably, we identified the complete chromosome of the candidate phylum *Candidatus Saccharibacteria*. Our results demonstrate that long-read based metagenomics provides a powerful tool for profiling plant-associated microbial communities.

## Materials and Methods

### Sampling and bacterial cell enrichment

Rice plants (*Oryza sativa* cultivar ‘Koshihikari’) were grown in an experimental paddy field at the Institute for Sustainable Agro–ecosystem Services, Graduate School of Agricultural and Life Sciences, The University of Tokyo (35°74′N, 139°54′E). Plants were sampled before heading on August 6^th^, 2018, and their roots and aerial parts were separated and stored at −80 °C. The aerial parts were ground using a Roche GM200 grinder (2,000 rpm 15 sec Hit mode, 8,000 rpm 30 sec Cut mode and 8,000 rpm 15 sec Cut mode) with dry ice. Plant-associated microbes were enriched from the ground samples using a bacterial cell enrichment method as previously described^19^.

### DNA extraction

Genomic DNA was extracted from the enriched plant-associated microbes using enzymatic lysis. The cells were lysed by the addition of 20 mg/ml Lysozyme (Sigma-Aldrich), 10 μl of Lysostaphin (>3,000 unit/ml, Sigma-Aldrich) and 10 μl of Mutanolysin (> 4,000 units/ml, Sigma-Aldrich), and incubated for 3 h at 37 °C. SDS (20%, Sigma-Aldrich) and Proteinase K (10mg/ml, Sigma-Aldrich) were then added, and DNA was purified with cetrimonium bromide and phenol-chloroform. DNA was then incubated with RNase for 30 min at 37 °C (Nippongene) and dissolved in TE buffer at 4 °C. The sequencing library was constructed and sequenced within a week of DNA extraction. We also extracted genomic DNA using a Fast DNA spin kit (MP-Biomedicals) for mechanical lysis from the enriched microbes. DNA fragmentation was assessed using pulsed-field gel electrophoresis.

### Sequencing of the V4 regions of 16S rRNA genes

The V4 regions of 16S rRNA genes were amplified using the primers 515F (5’-ACA CTC TTT CCC TAC ACG ACG CTC TTC CGA TCT GTG CCA GCM GCC GCG GTA A-3’) and 806R (5’-GTG ACT GGA GTT CAG ACG TGT GCT CTT CCG ATC TGG ACT ACH VGG GTW TCT AAT-3’) and sequenced with an Illumina MiSeq v3 platform. The first 20 bases of primer sequences were trimmed from all paired reads. For low-quality sequences, bases after 240 bp and 160 bp of forward and reverse primer sequences, respectively, were truncated using Qiime2 v2018.11.0^20^. The processed reads were aligned to the SILVA123 dataset^21^, and their taxonomy was provisionally determined.

### Sequencing, assembly, and gene annotation of the plant microbiome

SMRTbell libraries for sequencing were constructed according to the manufacturer’s protocol (Part Number 101-693-800 Version 01) without shearing. The libraries were cut off at 20 kbp using the BluePippin size selection system (Sage Sciences). Libraries were sequenced on two SMRT Cells 8M (Pacific Biosciences). We removed contaminating plant sequences, subreads showing more than 80% identity and 80% length coverage according to minimap2 v 2.14^22^ to ‘Nipponbare’ as the reference rice genome ^23^, and PacBio’s internal control sequences from subreads. ‘Nipponbare’ was used as the reference genome because the draft genome of ‘Koshihikari’ is highly fragmented, with an average read length of 32 bp^24^. The remaining subreads were assembled using Canu (version 1.8) ^25^ with the parameters previously described^26^. After assembly, we removed contaminated contigs derived from internal controls and reference genome using the same method, and artificial contigs with long stretches of G, C, A or T by calculation of GC contents with seqkit v0.11.0^27^. For quality assessment of the contigs^18^, we aligned the error-corrected reads generated during assembly to the contigs with pbmm2 v 1.2.1 (Pacific Biosciences) with maximum best alignment 1 and minimum concordance percentage 90 set as parameters, and extracted the contigs with a depth >5. Contig circularity was determined as previously described^26^. For confirmation of quality, error corrected reads were aligned to contigs using pbmm2 with the same parameters as for contig quality assessment, then gaps and coverage were assessed using IGV browser v 2.8.2^28^. Quast v 5.0.2^29^ was used to evaluate the quality of genome assemblies. Functional annotation of bacterial genes was conducted using PROKKA v 1.14.6 in the metagenomic mode^30^, COG database (BLASTP with the *e* value lower than 1e-05), Interproscan v 5.46-81.0 (with the *e* value lower than 1e-05) and kofamscan v1.3.0^31^. Augustus v. 3.4.0 was used to annotate the genes of fungal genomes^32^.

### Estimation of microbial composition using 16S rRNA genes

We obtained bacterial 16S rRNA gene sequences from NCBI BioProject 33175 (Bacteria) and 33317 (Archaea), removed those ≤ 1,400 bp in length, and clustered the remaining sequences using CD-HIT version 4.8.1 (≥ 98.7% identity) ^33^. The resulting curated 16S rRNA gene database contains 11,782 distinct sequences. In the long read-based assembly data, 16S rRNA genes longer than 1,400 bp on contigs having an average read depth ≥ 5 were aligned to our curated 16S rRNA gene database to obtain the maximum number of target hits. Alignments with <95% length coverage were removed. We used 16S rRNA genes with ≥ 98.7% identity as the top hits for approximating bacterial community composition at the species level. We counted the depth of the contigs carrying 16S rRNA genes to estimate their abundance.

Full-length 16S rRNA genes were amplified using the primers 27F (5’-AGR GTT YGA TYM TGG CTC AG) and 1492R (5’-RGY TAC CTT GTT ACG ACT T), and sequenced on a SMRT cell 1M v3. Circular consensus sequences (>3 paths) were constructed and demultiplexed using SMRTLink v 8.0.0 with default parameters. Primers and chimeric reads were removed from demultiplexed CCS reads using dada2 v 1.16^34^, and reads ≥ 1,400 bp were extracted. Full-length 16S rRNA amplicons were aligned with our curated database to assign taxa and to estimate bacterial community composition at the species level (≥ 98.7% identity). 16S rRNA gene sequences in the metagenomic data were aligned with MAFFT v7.475^35^ using default parameters. A phylogenetic tree was constructed using RAxML v8.2.12^36^ and visualized using FigTree v1.4.4 (http://tree.bio.ed.ac.uk/software/figtree/).

### Taxonomy assignment of large contigs

16S rRNA gene similarity and <u>a</u>verage <u>n</u>ucleotide identity (ANI) were used to assign contigs larger than 1Mbp (n = 172, including 6 circular contigs) to taxa with GTDB-tk v 1.1.1 using default parameters^37^. To provisionally assign taxa to contigs that were not assignable using either of the above methods, the annotated genes on the contigs were aligned to the nt database in NCBI using BLASTN with ≥ 80% identity and ≥ 80% length coverage. Comparisons of circular contig sequences to reference genomes were plotted with mummerplot, and contig completeness was calculated using checkM^38^. Circular contigs were classified as chromosomes or megaplasmids according to the criteria of diCenzo and Finan^39^. Interproscan was used to search for genes encoding replication proteins, such as DnaA and RepA.

### Classification of circular contigs less than 1 Mbp

The sequences of the circular contigs were compared to known sequences obtained from prokaryotic reference or representative genomes in the NCBI database and Plsdb version 2020_06_29^40^ using nucmer. Interproscan was used to search for plasmid-enriched or virus-related genes, and to classify those contigs as plasmid or phage. Kofamscan was used to annotate VirB/VirD4 systems on plasmids. Plasflow and Mob-typer via MOB-suite^41^ were also used for plasmid prediction, and for predictions of the plasmid host. Virsorter2 (a score cut off >0.8) ^42^ was used for predicting whether or not each contig originated in a bacteriophage. CheckV^43^ was used for assessing contig quality of viral sequences. To provisionally assign the taxonomy of each contig, we aligned all genes on the contig to the nt database in NCBI. The taxonomy of a contig was assigned as follows: if more than one-fourth of the genes on a contig showed > 80% identity and > 80% coverage to the corresponding genes of the nt database in NCBI, we assigned the taxonomy of the contig accordingly. If the genes on a contig were aligned to a strain of a genus, but to a different species, the taxonomy of the contig was estimated at the genus rank. Reliable taxonomy assignment was limited to cases where the number of the genes aligned to known sequences was greater than one-fourth of the contig, and the genes on the contig derived from one phylum. In all other cases, we concluded that the taxonomy of those contigs was unassigned. However, in some cases Mob-typer could assign the taxonomy of contigs which were not assigned using nt database. We tentatively assigned taxonomy using the Mob-typer result for these contigs. Gene maps were drawn with ‘ggplot2’ in R (https://www.R-project.org/).

### Taxonomic assignment and predicted gene function

We aligned all predicted genes to the COG database with an e value < 1e-05 to predict gene function^18^. A similarity search of the annotated genes was conducted against the nt database in NCBI using BLASTN with ≥ 95% identity and ≥ 90% coverage. From these genes, we extracted those which were identified by species, and counted both the number of genes and contigs that carried these genes.

### Genomic features of Candidatus Saccharibacteria (TM7)

We obtained the 16S rRNA genes of *Candidatus Saccharibacteria* (formerly known as TM7) from the NCBI database, removed sequences ≤ 1,400 bp, and clustered the remaining using CD-HIT at 98.7% identity. The genomic sequences of RAAC3 (GenBank: CP006915.1), *Candidatus Saccharimonas aalborgensis* (S_aal, GenBank: CP005957.1), GWC2 (GenBank: CP011211.1), TM7x (GenBank: CP007496.1) and YM_S32 (GenBank: CP025011.1) were obtained for genomic comparisons. Kofamscan and BlastKOALA^44^ were used to predict metabolic features. Average amino acid identity was calculated using the Kostas lab AAI calculator with default parameters (http://enve-omics.ce.gatech.edu/aai/).

## Data availability

Metagenomic data has been deposited in NCBI under accession number SAMN32580422 (BioSample), SRR23280466 and SRR23280465 (SRA) and PRJNA929667 (BioProject).

## Results

### DNA preparation for long-read metagenomics

To study rice-associated microbial communities in an agricultural environment, we harvested 8-week old rice plants from a paddy field and extracted their leaf-associated microbes using a density gradient centrifugation method to remove rice tissues^19^ (Extended Data Fig. 1A). We then compared enzymatic and mechanical cell lysis methods for their ability to extract intact genomic DNA with minimal damage using pulsed-field gel electrophoresis. This analysis showed that enzymatic lysis yielded intact chromosomes, whereas mechanically lysed cells yielded fragmented DNA (Extended Data Fig. 1B). Comparisons of bacterial community composition using the 16S rRNA gene V4 region indicated that the two methods gave similar results (Pearson’s correlation coefficient = 0.84, Extended Data Fig. 2), although some phyla were more highly represented in one method than in the other, possibly due to differences associated with cell lysis of different taxa. Given the importance of intact genomic DNA for long-read sequencing, we chose to use enzymatic lysis for further analyses.

**Figure 1.**
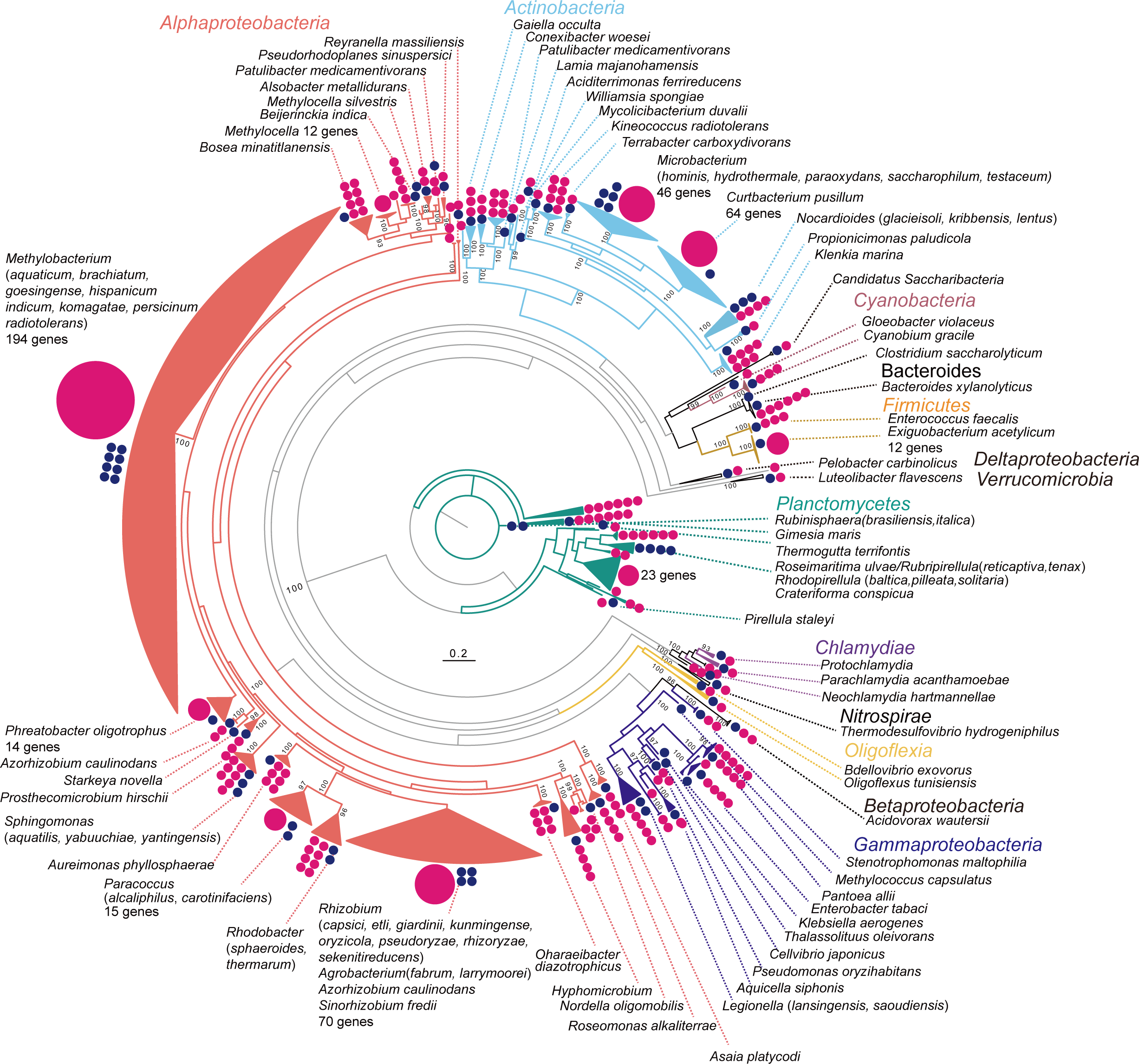
Overview of the phylogeny of 16S rRNA genes detected in the metagenome. Pink circles indicate the number of 16S rRNA genes detected in the metagenome for each major branch: large pink circles with numbers represent branches with more than ten genes, small pink circles each represent one gene. Blue circles represent the number of top species/genus identities of metagenome-derived 16S rRNA genes.

**Figure 2.**
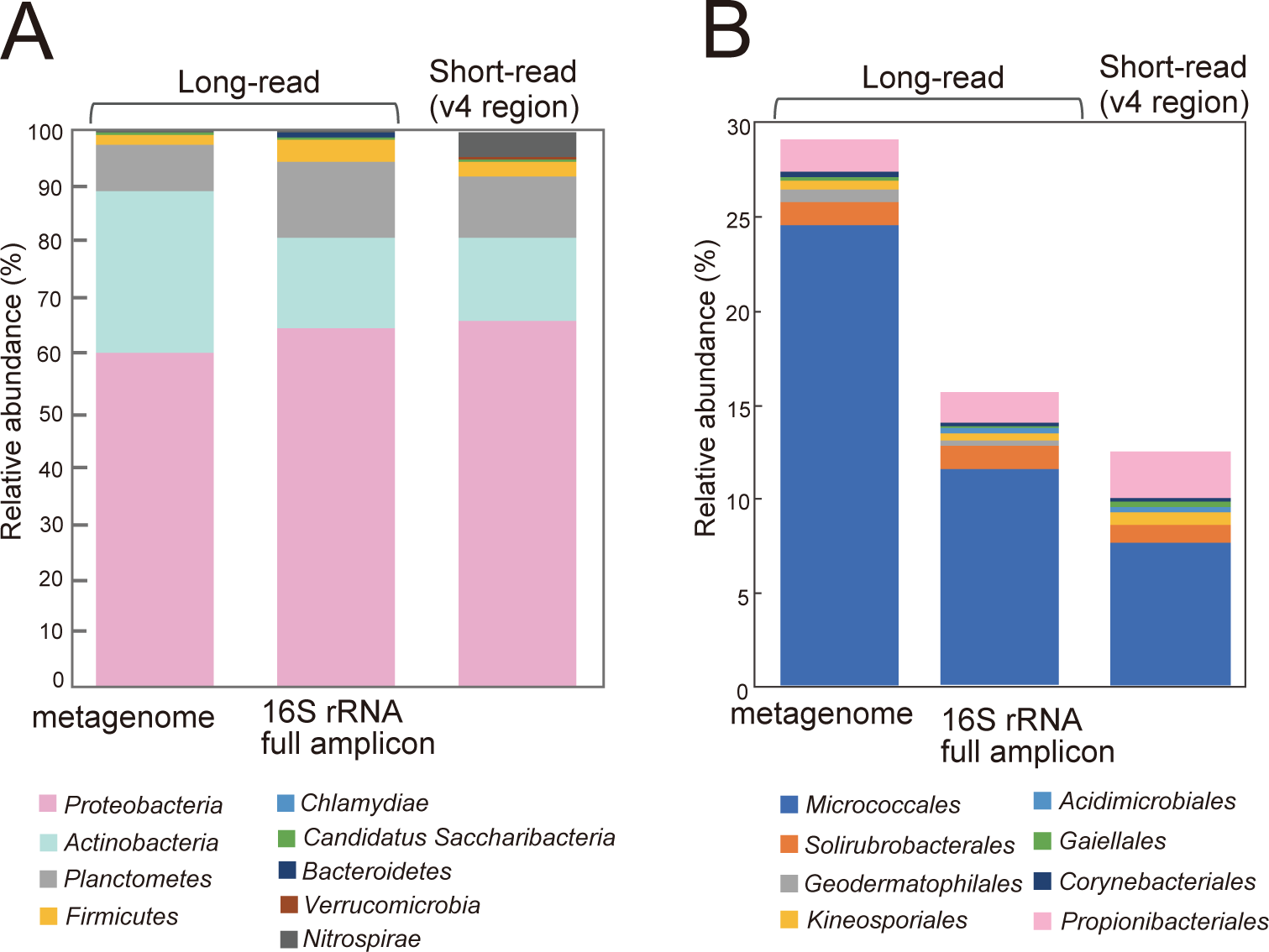
Relative abundance of 16S rRNA genes detected in the metagenome, compared to 16S rRNA gene full-length amplicon sequences and short reads. (A) Relative abundance of bacterial phyla and (B) relative abundance of the class *Actinobacteria* for each sequencing method.

### Long-read metagenomic sequencing of leaf-associated microbes

We sequenced gDNA from the leaf-associated microbes using two Sequel II 8M cells, yielding 140 Gbp of data with a mean read length of 17 kbp and a mean library size of 15 kbp (Supplementary Table1), indicating that our DNA extraction method was suitable for PacBio long-read sequencing. We obtained 26,067 contigs in total after assembly, with an N_50_ of 127 kbp, including 136 circular contigs smaller than 1 Mbp and 172 ≥ 1 Mbp, 6 of which were circularized (Table 1). A previous study reported that PacBio contigs with ≥ 1 and 5 read depths had ≥ 98.5% and 99.4% identity, respectively, when aligned to short-read contigs^18^. We thus were able to define 13,050 contigs with a depth of more than 5 as high quality contigs. These contigs represented approximately half of the total, and represented about 80% of the total reads (Table 1). Additionally, more than 90% of the total nucleotides were found in the set of contigs with a length of ≥ 50 kbp. Importantly, all large size contigs (≥ 1Mbp) were of high quality (Table 1). These data suggest that nucleotide sequences obtained from the rice phyllosphere microbiome are reliable for estimating bacterial community composition and their functions within the community.

**Table 1.**
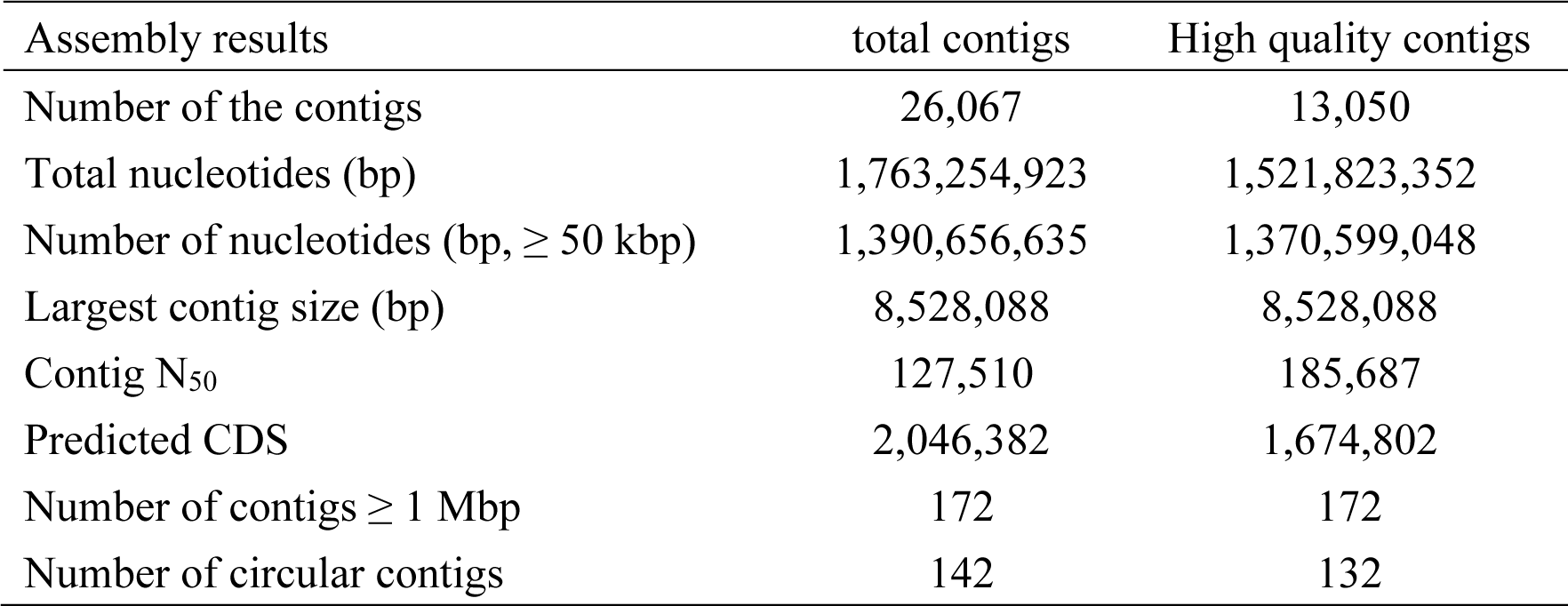
Summary of assembly results.

### Estimation of microbial composition using 16S rRNA genes in long-read metagenomics

We extracted 16S rRNA gene sequences ≥ 1,400 bp in length and ≥ 95% coverage of the top hits from high quality contigs from the metagenomic data. A total of 669 16S rRNA genes were identified on 561 contigs, representing 4.4% of high quality contigs (Supplementary Table 2). A phylogenetic tree was used to summarize the taxonomy of the detected 16S rRNA genes in the metagenome (Fig. 1). Many of 16S rRNA genes were clustered with top-hit sequences at various taxonomic ranks, such as the species and genus levels, but some were independently clustered. Of the 669 16S rRNA genes, 194 were clustered into the *Methylobacterium* genus. The 669 16S rRNA genes clustered into 166 bacterial species, 130 of which had ≤ 98.7% identity with any organism in the database, suggesting that they represent novel species (Supplementary Table 3). Clustering the 16S rRNA genes using a threshold for bacterial taxonomy^45^ showed that 290 of the 16S rRNA genes on 231 contigs were ≥ 98.7% identical to sequences attributable to known taxa (Table 2), but 378 were ≤ 98.7% identical to known taxa (Table 2). Among the latter, 16 were ≤ 82 and, 1 was < 78%, suggesting that they potentially represent a novel order and class, respectively (Table 2). Ten of the 16S rRNA genes represent a putative novel order within *Planctomycetes*.

**Table 2.** 16S rRNA genes above the threshold for bacterial taxonomy detected in the metagenome.

| Identity (%) | Taxonomic rank | number of 16S rRNA genes | number of contigs |
| --- | --- | --- | --- |
| 98.7 | species | 290 | 231 |
| 94.5 | genus | 234 | 198 |
| 86.5 | family | 127 | 122 |
| 82 | order | 16 | 16 |
| 78 | class | 1 | 1 |

Taxonomic analysis of the high-confidence identity 16S rRNA sequences (290 sequences with ≥ 98.7% identity) identified 40 bacterial species (Supplementary Table 4). For example, *Curtobacterium pusillum* was the most abundant species; 57 of 16S rRNA genes were identified on 46 contigs with a relative abundance of 15.2 %. Similarly, 8 species of *Methylobacterium* were identified in the high-confidence identity sequences, with three species (*M. indicum*, *M. radiotolerans* and *M. komagae*) having more than 10 16S rRNA genes (Supplementary Tables 2 and 4). The number of contigs containing 16S rRNA genes was mostly lower than the number of 16S rRNA genes, indicating that these bacteria carry multiple 16S rRNA gene copies (Supplementary Tables 2 and 4). In particular, nine of the 16S rRNA genes from *Exiguobacterium acetylicum* were detected on a single 1.7 Mbp contig (Contig ID: RRA86345 in Supplementary Table 2).

To verify bacterial community composition based on the long-read metagenome, full-length 16S rRNA genes were amplified using universal bacterial primers and sequenced to compare the taxonomic profiles. Among the 6,678 reads obtained, 2,958 reads (44%) have ≥ 98.7% identity to taxonomically classified 16S rRNAs, a result similar to the metagenomic data (Extended Data Fig. 3). We identified 64 taxa (Supplementary Table 4) to the species level, with *C. pusillum* being the most abundant, also consistent with the long-read metagenomic data. There were 16 species with relative abundance greater than 1%, 13 of which were also detected in the long-read metagenome (Supplementary Table 4). The combined relative abundance of the 13 species was 42.0% in the metagenome and 33.3% in the 16S rRNA nearly full-length amplicon data sets.

We also compared the relative abundance of the rice bacterial community determined by long-read metagenome sequencing versus short-read 16S rRNA sequencing (Fig. 2), and found that the relative abundance of *Actinobacteria* was about twice as high in the long-read metagenome (28.6%) than in either the nearly full-length 16S rRNA amplicon dataset (15.8%) or the V4 region dataset (15.2%). Comparing the relative abundance of *Actinobacteria* showed that the relative abundance of *Micrococcales* in the metagenome was about twice that of PCR-based amplicon sequencing. The difference may be because certain classes of *Actinobacteria* were difficult to detect using universal primers, even though the target sequences are identical^46^. Previous studies also showed that the V4 region is less reliable for classifying *Actinobacteria* sequences^47^. Therefore, our results suggest that long-read metagenome sequencing provide more accurate identification about dominant bacterial communities in the aerial parts of rice, particularly for *Actinobacteria*.

### Taxonomic assignment of predicted genes

We identified a total of 2,046,382 predicted genes in the metagenome (Table 1, Extended Data Fig. 4). Of these putative genes, 364,262 had an e-value of 1.0e-05 or less and were annotated using the COG database. Approximately 20% of the genes were categorized as poorly characterized group ‘R’ (11.8%, general function prediction only) or ‘S’ (9.6%, function unknown). 8% of the genes were annotated as amino acid metabolism (E), 6.5% as carbohydrate transport and metabolism (G), and 6.2% as energy production and conversion (C). Among these five categories, 50 - 70% of the genes were derived from *Alphaproteobacteria*, particularly *Methylobacterium* (12.9 - 17.3 %). The putative genes from Planctomycetes were the second most abundant (9.6 – 18.0 %, Extended Data Fig. 4). These results showed that *Methylobacterium* is the dominant genus in the rice phyllosphere, supporting the bacterial community composition predicted by 16S rRNA genes (Fig. 1).

### Taxonomic assignment of large contigs

16S rRNA gene sequences and ANI (GTDB-tk) were used to assign the taxonomy of 172 high quality contigs, which ranged from 1 Mbp to 8.5 Mbp (Fig. 3, Supplementary Table 5). 16S rRNA genes were found in 98 contigs (97 chromosomal and one in a megaplasmid), and ANI was able to assign taxonomy of 147 of the contigs (Fig. 3, Supplementary Table 5). 92 contigs were assigned by both the 16S rRNA gene and ANI.

**Figure 3.**
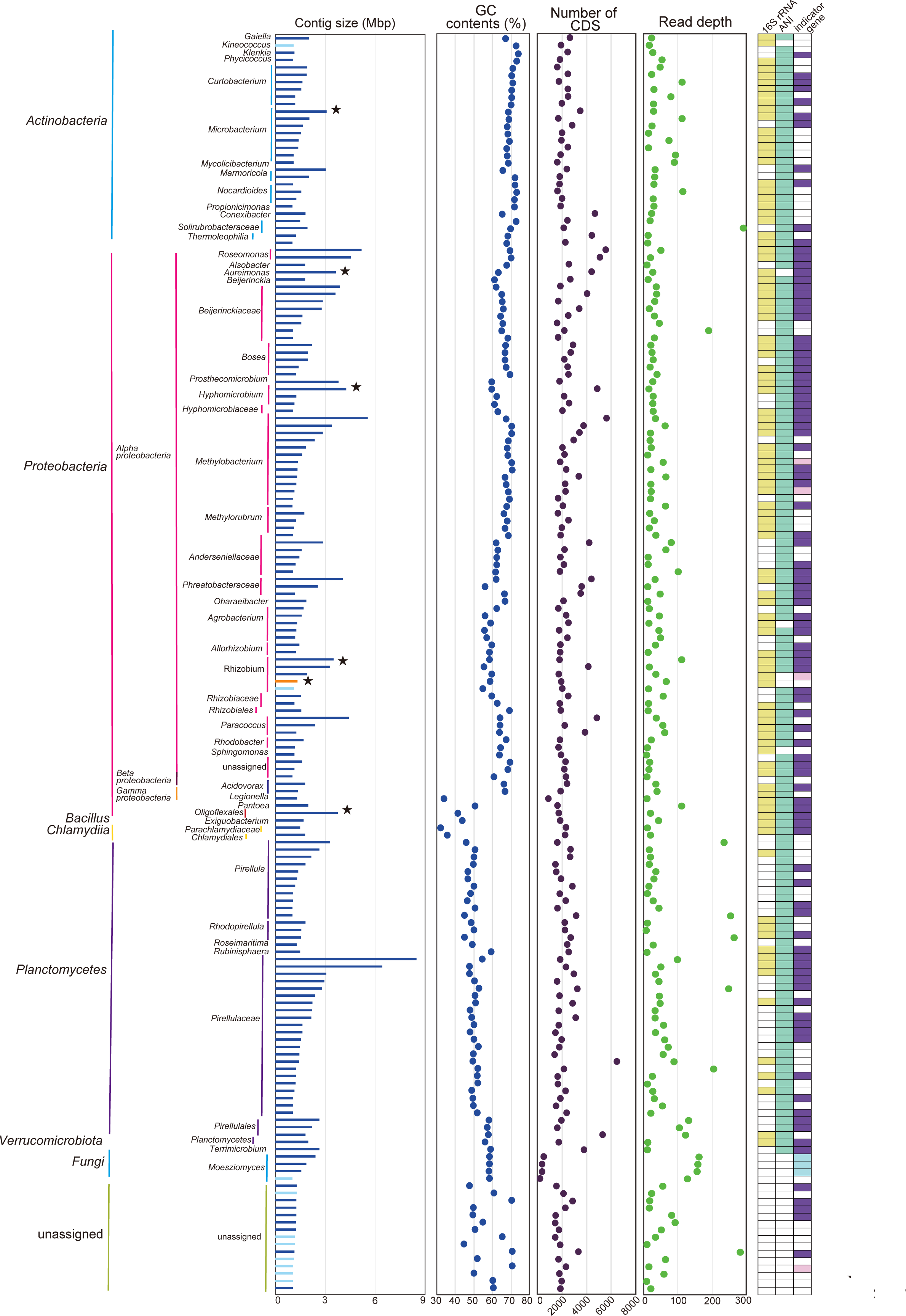
Characteristics of large contigs (>1Mbp, n=172). The taxonomic assignment of each contig was determined based on 16S rRNA genes, ANI, and gene similarity searches. Contigs carrying 16S rRNA genes are shown in yellow blocks. Contigs classified by ANI are shown in green blocks. Contigs carrying *dnaA*, *repA* and mini-chromosome maintenance genes are shown in purple, pink and light blue blocks, respectively. The blue, red and black bars in the contig size column represent chromosomes, megaplasmids and unclassified (neither chromosome nor plasmid), respectively. The black star indicates a circular contig.

We also assigned the taxonomy of one contig using a similarity search of its genes. In total, the taxonomy of 157 contigs was assigned by these methods (Fig. 3, Supplementary Table 5), but 19 contigs could not be assigned using either method. Among these, the genes of 4 contigs showed a high identity (≥ 95%) and length coverage (≥ 90%) to the *Moesziomyces antarcticus* (Fig. 3, Supplementary Table 5), suggesting that these four contigs originated from a yeast.

We also attempted to discriminate between chromosomal and plasmid contigs using ANI and the presence or absence of DNA replication initiators such as DnaA (for bacterial chromosomes) and RepA (for some plasmids). We classified only one contig as a megaplasmid (RRA6539; Fig. 3, orange color in the contig size column), as it showed high similarity to a plasmid (NZ_CP049244.1) of *Rhizobium pseudoryzae*, which also carries 16S rRNA genes on both its chromosome and plasmid. In addition, four contigs were derived from the yeast *M. antarcticus* using blast search and three of four contigs were carried minichromosome maintenance proteins (MCM2, 3, 6 and 10), suggesting that those three contigs may be a minichromosome of *M. antarcticus* (Fig. 3, Supplementary Table 5). In total, 156 of the large high quality contigs were classified as bacterial chromosome, one was a megaplasmid, and four were fungal sequences. The other 11 large contigs were not classified as either chromosomal or plasmids using these methods.

Of the 156 bacterial chromosomal contigs, five were circularized, suggesting that they are complete chromosomes (Black star in Fig. 3, Supplementary Table 5). The taxonomy of these genomes can be tentatively assumed by 16S rRNA gene sequence similarity and/or ANI, though the nucleotide sequences of some contigs do not match those of sequenced strains (Extended Data Fig. 5). For example, four contigs (RRA2267, RRA3045, RRA85519 and RRA944769) carry 16S rRNA genes with 87.4% - 99% identity to the top hit (Fig. 3 and Supplementary Table 5). In particular, RRA944769 could be classified as a complete genome of a novel family of *Oligoflexales* based on 16S rRNA gene identity and ANI (Fig. 3 and Supplementary Table 5). The other three contigs (RRA2267, RRA3045 and RRA85519) and one additional contig (RRA944769) were placed at the genus and order rank by ANI, respectively, all of which were consistent with the 16S rRNA-based assessment (Fig. 3 and Supplementary Table 5). Curiously, no 16S rRNA gene was detected in RRA2326, but it was assigned to the genus *Aureimonas* by ANI (Fig. 3 and Supplementary Table 5). This confirms a previous report showing that *Aureimonas* sp. AU20, isolated from the rice phyllosphere, has its 16S rRNA gene on a small plasmid, but not on the chromosome^48^. Additionally, the 11 unclassified contigs were shown to have >90% completeness and <5% contamination by CheckM, suggesting that those 11 contigs were nearly-complete chromosomes^38^.

### Classification of circular contigs smaller than 1Mbp

Of the 136 circular contigs ranging in size from 8.5 kbp to 832 kbp, with the GC content from 36.8% to 75.2% (Fig 4, Supplementary Table 6), the sequences of 134 did not align to any known sequences, suggesting that these are novel sequences. The remaining two contigs were aligned with high similarity to a plasmid of *Methylobacterium phyllosphaerae* strain CBMB27 (NZ_CP015369.1) (red color in contig size column in Fig 4, Extended Data Fig. 6). 61 genes on 41 of the 136 contigs were annotated as *repC* (Fig 4, Supplementary Table 6), suggesting that these contigs are novel *repABC* plasmids. Plasmid hosts were identified using a similarity search from construction of a phylogenetic tree using the RepC protein sequences and mob-typer. Sixteen contigs were associated with *Alphaproteobacteria*, while the remaining 23 contigs could not be assigned using these methods (Fig 4, Extended Data Fig. 7, Supplementary Table 6). These results show that the likely origin of nearly two-thirds of the *repABC* plasmids detected in this study is bacteria that have not been reported to carry *repABC* plasmids.

**Figure 4.**
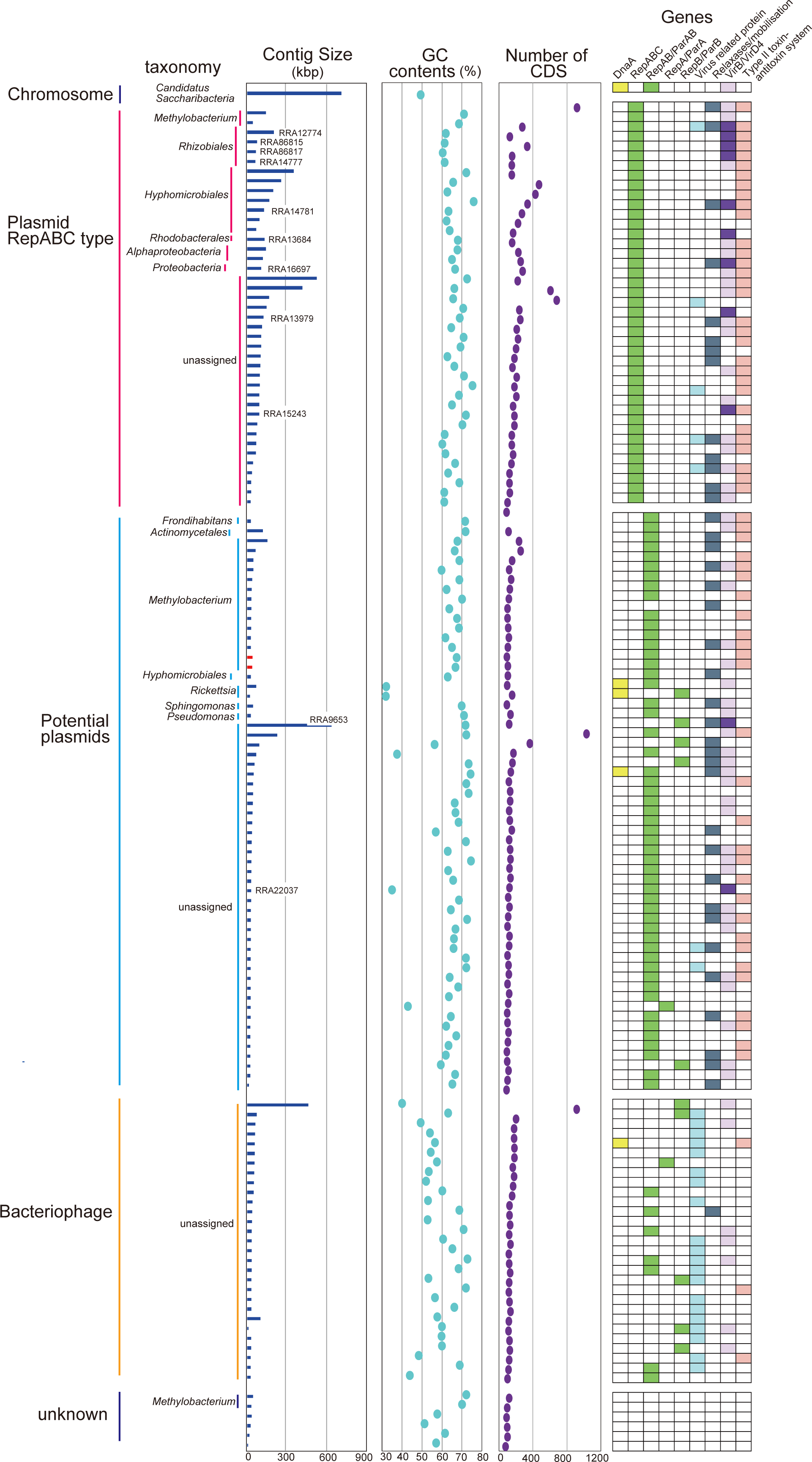
Characteristics of small circular contigs (<1Mbp, n=136). Gene annotations are indicated by color blocks. Dark purple and light purple represent complete/nearly complete, and partial genes of VirB/VirD T4SS systems, respectively. Red bars in the contig size column indicate known plasmid sequences. The contig ID shown in the contig size column corresponds to contigs carrying complete/nearly complete VirB/VirD T4SS.

An additional 29 contigs were classified as dsDNA bacteriophages with a high score (> 0.8) from virsorter2 with a CheckV completeness ranging from 14.3 to 100% (Fig 4, Supplementary Table 6). Twenty-one of these contigs carried putative phage-related genes, such as for capsid proteins. Thirteen carried putative partitioning protein genes, seven contigs carried VirB/VirD4 component genes, and one had a relaxosome protein TraY gene (Fig 4, Supplementary Table 6). Although we identified presumptive novel bacteriophages in this study, we were unable to assign their taxonomy.

We identified 59 contigs that carried presumptive VirB/VirD4 component, relaxosome protein, or type II toxin-antitoxin system genes, but were not classified as either plasmids or bacteriophages by mob-typer and virsorter2. These genes are more commonly plasmid-borne than chromosomal^5^, suggesting that these contigs may represent incomplete plasmids (Fig 4, Supplementary Table 6). Similarity searches of the genes on these contigs suggested that 21 out of the 59 contigs were from *Alphaproteobacteria*, *Gammaproteobacteria* (*Pseudomonas*), or *Actinobacteria* (Fig 4, Supplementary Table 6).

Pathogenic bacteria inject their Type 4 Secretion System (T4SS) effector molecules directly into host cells, thereby altering host cell functions. The 11 gene products of the *Agrobacterium tumefaciens virB* operon, together with the VirD4 protein, are thought to form a membrane complex which facilitates the transfer of T-DNA to plant cells. VirB/VirD4 T4SS components were present on 11 contigs (Fig. 5), nine of which were *repABC* type plasmids, and the remaining two were possibly plasmids. The likely origins of seven of these contigs were *Proteobacteria*, including *Rhizobiaceae*, and *Rhodobacterales* in the *Alphaproteobacteria* group. The remaining four could not be assigned to a taxonomic group. A comparison of the gene arrangements on the 11 contigs with those of *A. tumefaciens* showed that all, or almost all components of VirB/VirD4 T4SS were present, although in a different order than in *A. tumefaciens*, with some gene duplication and missing components. An additional 52 contigs carried at least one component gene of the VirB/VirD4 T4SS.

**Figure 5.**
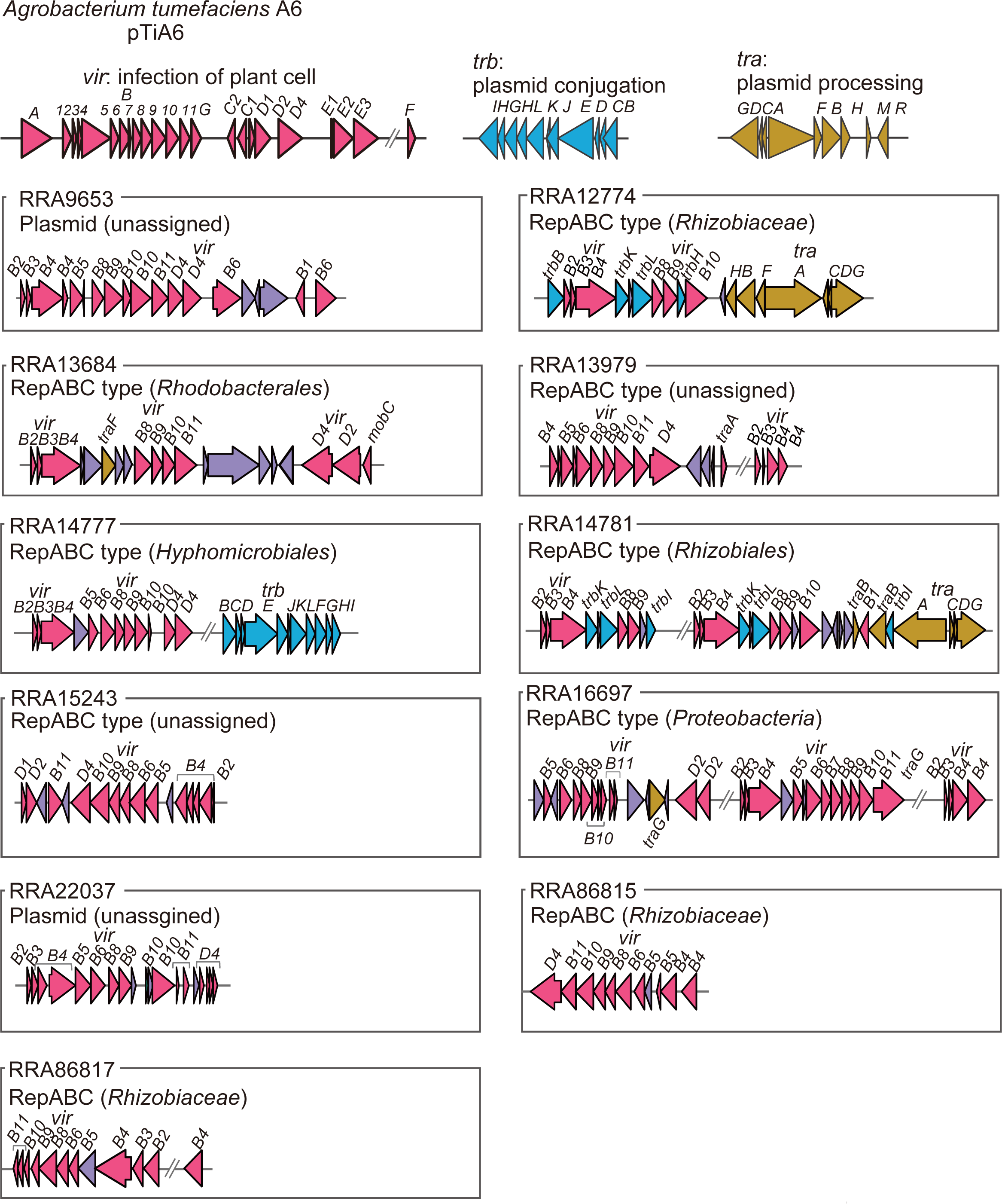
Gene arrangements, predicted host, and estimated type of plasmid for T4SS genes discovered in the metagenome. Purple indicates hypothetical or non-T4SS component genes.

We identified five small circular contigs carrying a presumptive *dnaA* gene, which is typically found in bacterial chromosomes as part of the DNA replication machinery (Fig. 4 and Supplementary Table 6). A similarity search revealed that the origin of two of those contigs was *Rickettsia*, which are obligate intracellular *α-proteobacteria* associated with various eukaryotic hosts. Notably, approximately half of the 26 validated *Rickettsia* species have plasmids, some of which carry a *dnaA*-like gene and range from in size from 12 kb to 83 kb^49^. We also detected *dnaA* genes on contigs that were classified as bacteriophage, potentially plasmid, or from the *Candidatus Saccharibacteria* chromosome. Two other contigs originated from *Methylobacterium,* but we were unable to classify these contigs as plasmid or bacteriophage based on the available gene information. Four contigs could not be classified as chromosome, plasmid, or bacteriophage due to a lack of similarity to any known bacterial or bacteriophage-derived genes in public databases. In total, our analysis presumptively identified one chromosome, 100 plasmids (41 *repABC*-type plasmids and 59 potentially plasmid-associated contigs), 29 bacteriophages, and six unclassified contigs, demonstrating that long-read metagenomic sequencing can effectively be used to identify a large number of plasmids from a complex microbial community, most of which are novel.

### Complete genome of a bacterium in the *Candidatus Saccharibacteria* as-yet uncultured phylum

One of the key benefits of long-read metagenomic analysis is the potential to obtain complete genome sequences of uncultivable microorganisms. Here, we obtained the whole chromosome sequence of a member of the uncultured *Candidatus Saccharibacteria* phylum as a circular contig (RRA8490). Phylogenetic analysis based on 16S rRNA genes indicated that RRA8490 clusters with isolates found in the human oral microflora (Extended Data Fig. 8A). A comparison of whole genome sequences and amino acid identities between RRA8490 and previously determined strains in the *Saccharibacteria*^50–54^ showed that the genomes of these strains and RRA8490 were distinct, with average amino acid identities ranging from 52.2 to 54.2% (Extended Data Fig. 8B and C). Using Kofamscan and Interproscan, we searched for metabolism-related genes in RRA8490 and found that it encoded all the presumptive genes necessary for the biosynthesis of peptidoglycan (MurABCDEFG, MraY and MtgA), suggesting that its cell wall is of the gram-negative type (Fig. 6). Unlike other strains, RRA8490 did not encode amino acid or fatty acids synthesis genes^55^, but it did presumptively encode enzymes that metabolize glucose to ribose and glycerate-3-phosphate, as well as phosphoenol-pyruvate to malate, suggesting that these pathways may be used to generate ATP. RRA8490 also encoded four regions of type IV pili (*pilM_1_N_1_O_1_B_1_TC_1_D*, *pilB_2_C_2_M_2_N_2_O_2_*, *pilB_3,_ pilB_4_*), similar to a previously reported *Saccharibacteria* (TM7) genome that carries type IV pili for host cell attachment^53^ (Fig. 6). In addition to these characteristics, RRA8490 also encoded the cytochrome oxidase complex CyoABCDE, as previously reported.

**Figure 6.**
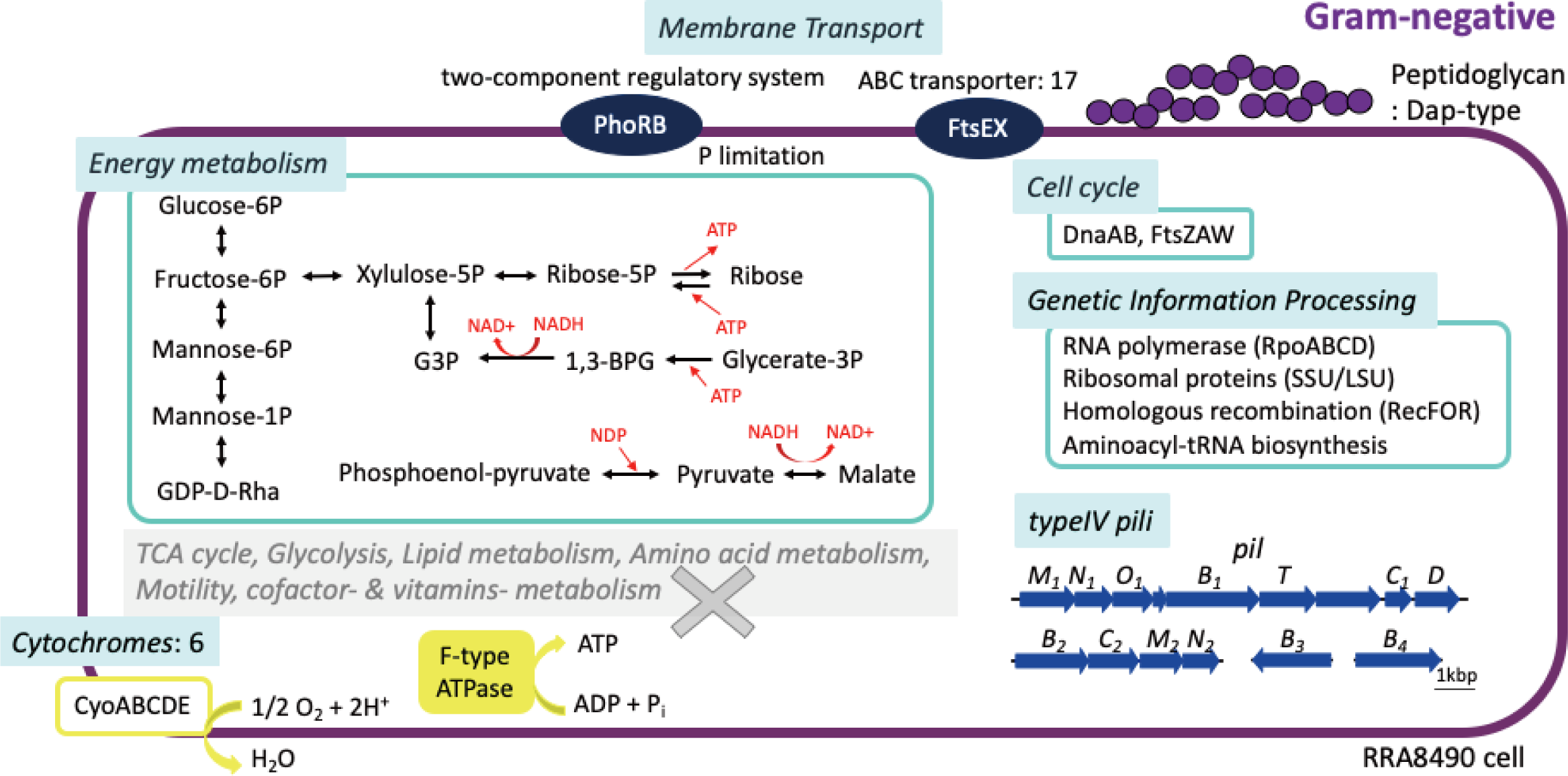
Predicted metabolism of RRA8490, a potential new strain in the *Candidatus Saccharibacteria* genus. Metabolic pathways were reconstructed using kofamscan, Interproscan and similarity searches.

## Discussion

This study demonstrated that enzymatic genomic DNA extraction combined with long-read metagenomic sequencing is an effective tool for profiling plant microbiota and their genomes, as well as defining complete chromosomes, plasmids and bacteriophages from long, high quality contigs. Importantly, comparisons of the community-profiling datasets obtained using short-read 16S rRNA amplicon sequencing confirms that the enzymatic DNA extraction method is largely unbiased, with the notable exception of the inclusion of fungal DNA from the *Moesziomyces* genus. This is not surprising, as *Moesziomyces* spp. are commonly detected in plants^56, 57^, but not amplified by bacterial 16S primers. Therefore, the community profile of rice leaves obtained using long-read metagenomics is consistent with previously reported datasets. For example, our data indicate that *C. pusillum* is the dominant species in rice leaves (Supplementary Table 2 and 4), which is consistent with the fact that *Curtobacterium* spp. have been isolated from the leaves of many different plants^58–61^ and are known to be abundant in a leaf litter communities^62^. Similarly, the *Methylobacteriaceae* family is a dominant presence (Fig. 1), as seen in the aerial parts of many plants^63^.

A potential 130 novel species were identified from 669 16S rRNA genes. The relative abundance of 16S rRNA genes detected in the metagenome ranged from 0.02% to 1.8%, demonstrating the depth of coverage provided by long-read metagenomics. However, a comparison between bacterial species detected in the metagenome and nearly full-length 16S rRNA amplicon sequencing revealed that some species were only detected using one method or the other, despite having relative abundances greater than 0.02% (Supplementary Table 4). This highlights the importance of using a combination of long-read sequences, such as metagenomes, and 16S rRNA amplicon sequencing for more comprehensive taxonomic assignments. Overall, long-read metagenomics is a powerful tool for accurately identifying bacteria in the rice phyllosphere, with the potential for greater discrimination between organisms as new analysis methods become available.

The use of long-read metagenomics allows for the estimation of bacterial species abundance currently possible, as it is not influenced by PCR amplification bias and variations in the number of 16S rRNA genes present in each genome. Indeed, the prevalence of *Micrococcales* (*Actinobacteria*) identified from metagenomic data was about two-fold higher than from amplicon data (Fig. 2). The number and location of rRNA operons (*rrn*) can vary significantly among bacteria, and some bacteria have multiple copies of *rrn* on different chromosomes, as seen in *Brucella*^64^ and *Vibrio*^65^. Our data demonstrated that the relative abundance of *E. acetylicum* is often overestimated due to the presence of nine copies of 16S rRNA genes on four contigs in the draft genome of *E. acetylicum*^66^. In contrast, long-read metagenomics-based identification of the precise number of 16S rRNA genes allows for accurate determination of bacteria abundance within a community. When 16S rRNA genes were not present on contigs, we were able to use ANI to tentatively assign taxonomy for nearly 30% of the contigs (Fig. 3 and Supplementary Table 5), because long-read metagenomics allows the reconstruction of very large contigs.

The use of long reads allowed us to identify the genes contained within six large circular contigs. Two of the contigs (RRA2326 and RRA3045) appeared to be derived from known species, but the others had little similarity to previously whole-genome sequenced strains (Fig. 3 and Supplementary Table 5), suggesting that the four remaining circular contigs represent the chromosomes and a megaplasmid of novel species. Notably, the complete chromosome of *Rhizobium giardini* (RRA85519) was sequenced and identified for the first time, which can serve as a valuable reference for this species. A novel strain in the genus *Oligoflexia* (RRA944769) was also identified, which differed in genomic composition and presumptive energy metabolism pathways from previously isolated and sequenced strains^67, 68^. For instance, RRA944769 encodes putative nitrate-and nitrite reductases (NO_3_ to NO), whereas *Oligoflexia tunisience* Shr3^T^ has genes converting from NO_2_ to N_2_. *O. tunisience* Shr3^T^ also has *aa*_3_-and *cbb*_3_-type cytochrome c oxidases, whereas RRA944769 has cytochrome *b*_6_ in addition to *aa*_3_-and *cbb*_3_. Our study also confirmed the natural occurrence of a chromosome lacking 16S rRNA genes in an *Aureimonas* sp. genome (RRA3045), resolving previous uncertainty around chromosomal rearrangements during cultivation^48^. These examples demonstrate the power of long-read metagenomics in accurately identifying and characterizing the genetic makeups of a complex sample.

Long-read metagenomics has a major advantage over short-read metagenomics in that it can be used to define circular mobile elements such as plasmids and temperate bacteriophages. These elements play a significant role in microbiome interactions and horizontal gene transfer^8, 69^. Short-read methods have largely been unable to fully assemble complete plasmids and bacteriophages, meaning that many of the genes present on these elements have not been identified. In this study, among 136 small circular contigs, only two were found to align well with known plasmids of *M. phyllosphaerae*, suggesting that the majority of these sequences represent novel plasmids or bacteriophages. Contigs that encode functions often found on plasmids, such as toxin-antitoxin systems and T4SS, are likely to be plasmids^18^. Those functions are often important for modulating interactions with plant cells and other microorganisms^70^. Approximately 22% of these contigs putatively encode RepC, a protein involved in *repABC* plasmid replication, which are common in *Alphaproteobacteria*^71^. In fact, almost half of the RepC-encoding genes identified in the novel plasmids appear to be clustered with *Rhodobacteraceae*, *Rhizobiales* and *Hyphomicrobiales*. Other RepC-encoding genes could not be assigned to any taxon (Fig. 4 and Extended Data Fig. 7), suggesting that they originated from unidentified bacteria. However, predicting a plasmid host, particularly for broad host range plasmids, is a challenging task in itself, let alone for metagenomic studies^9^. While we have made suggestions about the host of origin for some of the plasmids, the origin of others remains unclear. New technologies, such as droplet microfluidics for isolation of single bacterial cells combined with plasmid-specific markers may help to address this deficiency in the future^72^.

One of the best studied plasmid-specific functions is the VirB/VirD4 system of the Ti plasmid from *Agrobacterium tumefaciens*. The *virB* operon (*B1-11*) together with *virD4* encode a putative T4SS. T4SS are highly diverse^73–77^, and the exact number of genes and their role in T4SS assembly or function is unknown in many classes of T4SS^78^. The VirB/VirD4 system identified in RRA9653 carried *virB1-11* and *virD4* gene homologs (Fig. 5). Four additional putative plasmids (RRA13979, RRA16697, RRA22037 and RRA86817) were conserved for *virB2*-*B11* of the eleven *virB* and *virD4* genes, which are essential for construction of the VirB/VirD4 system in *A. tumefaciens* (Fig. 5), suggesting that these five plasmids are the VirB/VirD4 system of *A. tumefaciens* type. Interestingly, Ti plasmids belong to the *repABC* family, which is widely distributed among many species of *Alphaproteobacteria*. However, no *repABC* genes were found on two of the plasmids (RRA9653 and RRA22037), suggesting that these plasmids may have been horizontally transferred from other bacterial taxa.

Long-read metagenomics provide genomic information for poorly studied or as-yet uncultivated bacteria. We analyzed the genes identified in the metagenomic data that occurred with high confidence identity and coverage to specific bacterial species, and counted the number of contigs carrying these genes. Interestingly, the number of genes from *Planctomycetes bacterium* was much lower than expected, at 1,497 genes, despite being carried by the largest number (405) of contigs (Extended Data Fig. 9). Similarly, the number of the genes from *Microbacterium testaceum* and *Phreatobacter cathodiphilus* was also low, but these genes were carried by a large number of contigs. These findings suggest that the genomic information of these three species is largely unknown. For instance, our study found a high abundance of *Planctomycetes*: 56 out of 669 16S rRNA genes and 42 out of 172 large size contigs were derived from novel *Planctomycetes* (Figs. 1 and 3, Supplementary Tables 2 and 5). Because *Planctomycetes* are difficult to culture^79^, there is limited gene/genomic information available for this group. To our knowledge, the complete genome sequence of *M. testaceum* has only been deposited for one strain (3.98 Mbp) ^80^, but using long-read metagenomics, we were able to obtain seven large contigs that represent complete or nearly complete chromosomes of *Microbacterium* (Fig. 3). We were able to obtain 79 contigs from *P. cathodiphilus*, of which only one strain has been sequenced^81^ (Extended Data Fig. 9). Thus, our methodology can be used to parse the ecological and biological functions of fastidious bacterial groups.

Our long-read metagenomic analysis was able to define the complete circular genome (RRA8490) of an uncultured bacterium belonging to the *Candidatus Saccharibacteria* phylum. Members of this phylum have been detected in numerous natural environments such as soils, animals, and plants, but lack of cultured isolates has limited our understanding of their biology. Consequently, only a few complete genomes from this phylum have been reported^50–54^. Compared to the recently nearly completed genome (1.45 Mb) of an oat-associated member of the *Candidatus Saccharibacteria* phylum tentatively designated *Teamsevenus rhizospherense* strain YM_S32, RRA8490 is much smaller (0.83 Mb) and belongs to a different clade (Extended Data Fig. 8). Both *T. rhizospherense* and RRA8490 apparently lack the ability to synthesize amino acids from central metabolites, but RRA8490 is predicted to carry type IV pili and cytochrome bo3, similar to others in *Candidatus Saccharibacteria* phylum. RRA8490 is predicted to be able to assimilate and metabolize glucose and fructose, which are compounds found in leaf exudates of the rice phyllosphere^82^. This suggest that RRA8490 may utilize these compounds as carbon sources. Additionally, *Candidatus Saccharibacteria* are obligate epibionts of *Actinobacteria*, which they lyse to obtain nutrients^53^, suggesting that RRA8490 may not rely solely on plant exudates for nutrient acquisition, but may also degrade *Actinobacteria*. The CyoABCDE, cytochrome o oxidase complex, is used by *Rhizobium etli* to adapt to anaerobic conditions^83^, but Cyo appears to be produced only under oxygen-rich growth conditions in *E. coli*^84^. These results suggest that the ability to function at a wide range of oxygen concentrations, as demonstrated by the presence of Cyo in RRA8490, would be beneficial for this bacterium as it adapts to a variety of oxygen conditions in its natural environment.

In conclusion, long-read metagenomics fueled by high quality DNA extraction provides an efficient method for exploring uncharted organisms in the plant microbiome, and the resulting data represents an emerging primary resource for a deeper understanding of plant-associated microbial ecology.

## Acknowledgements

We are grateful to the technical staffs of the Department of Technical Development in ISAS of The University Tokyo for their maintenance of plant materials. We also thank Prof. Dr. K. Minamisawa and Dr. S. Hara for sharing the cell-density centrifugation from rice plants protocol. This work was supported by the JSPS KAKENHI grants JP20H05909 and JP22H00364 (K.Sh.) and JP20H05592 (S.M.) and by JPNP18016 commissioned by the New Energy and Industrial Technology Development Organization (NEDO).

## Contributions

All authors substantially contributed to this work. All authors approved the submitted version of this manuscript. Data analysis and interpretation of data were performed by all authors. S.M., and K.Sh. contributed to the design of the work. S.M., P. G., K. Sa and K. Sh carried out rice sampling from an experimental field plot.

## Extended Data

Extended Data Fig. 1. Preparation of genomic DNA from the rice-microbiome for long-read metagenomic sequencing. (A) Rice plants were sampled from experimental field and ground with dry ice. Bacterial cells were purified from aerial parts of rice plants using cell density centrifugation. (B) Genomic DNA was extracted from the purified microbiome using physical and enzymatic lysis. The presence of chromosomal DNA was confirmed using Pulse-field gel electrophoresis.

Extended Data Fig. 2. Comparison of the relative abundance of 16S rRNA genes extracted with mechanical lysis and enzymatic lysis.

Extended Data Fig. 3. Comparison of the relative abundance of 16S rRNA genes at different taxonomical ranks in the metagenome and 16S rRNA full-length amplicon sequences.

Extended Data Fig. 4. Gene categorization using the COG database. The number of genes categorized in each group is indicated in the heatmap.

Extended Data Fig. 5. Alignment of whole genomic sequences between the six circular contigs and the closest bacterial relative genome. The genome of the reference strain, *Rhizobium glardinii*, was determined by whole genome sequencing (WGS). The bold dotted line of the horizontal axis represents each contig of *R. giardinii*.

Extended Data Fig. 6. Alignment of the whole genomic sequences of RRA17620 and RRA19473 to the plasmid of *Methylobacterium phyllosphaerae* strain CBMB27 (NZ_CP015369.1).

Extended Data Fig. 7. RAxML phylogenetic tree of RepC protein encoded in small circular contigs (< 1Mbp). The 61 copies of RepC on 39 contigs were used to construct the tree. The accession number indicates the representative RepC in each cluster. The taxonomic assignment of RepC copies that are independently clustered with known RepC are defined as unknown. *Klebsiella pneumoniae* RepC was used as the outgroup.

Extended Data Fig. 8. Comparison of the *Candidatus Saccharibacteria* strain located in the long read metagenomic dataset with published genomes of other strains in the genus. (A) Phylogenetic tree using 16S rRNA genes of strains in *Candidatus Saccharibacteria*. Contig RRA8490 is indicated by the red box, adjacent to the human oral cavity group. *Candidatus Gracilibacteria* and *Candidatus Absconditabacteria* were used as outgroups. (B) Comparison of whole genomic sequences between RRA8490 and five *Candidatus Saccharibacteria* (TM7) isolates. Whole genomic sequences were compared using nucmer, showing that RRA8490 is not similar to the others. (C) Amino acid identity between RRA8490 and five isolates of *Candidatus Saccharibacteria*. The average amino acid identity was calculated using the AAI calculator developed by the Kostas lab with default parameters (http://enve-omics.ce.gatech.edu/aai/).

Extended Data Fig. 9. The number of genes identified in the metagenome dataset with high identity and coverage to specific bacterial species and of the contigs carrying these genes. The contigs with a minimum of 50 predicted genes are shown.

## Supplementary tables

Table 1. Summary of the sequencing results in this study.

Table 2. Summary of 16S rRNA genes detected in the metagenome dataset.

Table 3. Summary of the 16S rRNA genes clustered with ≥ 98.7% identity in metagenome dataset.

Table 4. All 16S rRNA genes (≥ 98.7 identity) detected in the metagenome, and full-length 16S rRNA gene amplicon sequences.

Table 5. Summary of large contigs (>1Mbp, n=172)

Table 6. Summary of small circular contigs (<1Mbp, n=136)

## Notes

### Competing Interest Statement

The authors have declared no competing interest.

