## Extended Data Fig for "Uncovering plant microbiomes using long-read metagenomic sequencing": 230222_ExtendedFigures.pdf

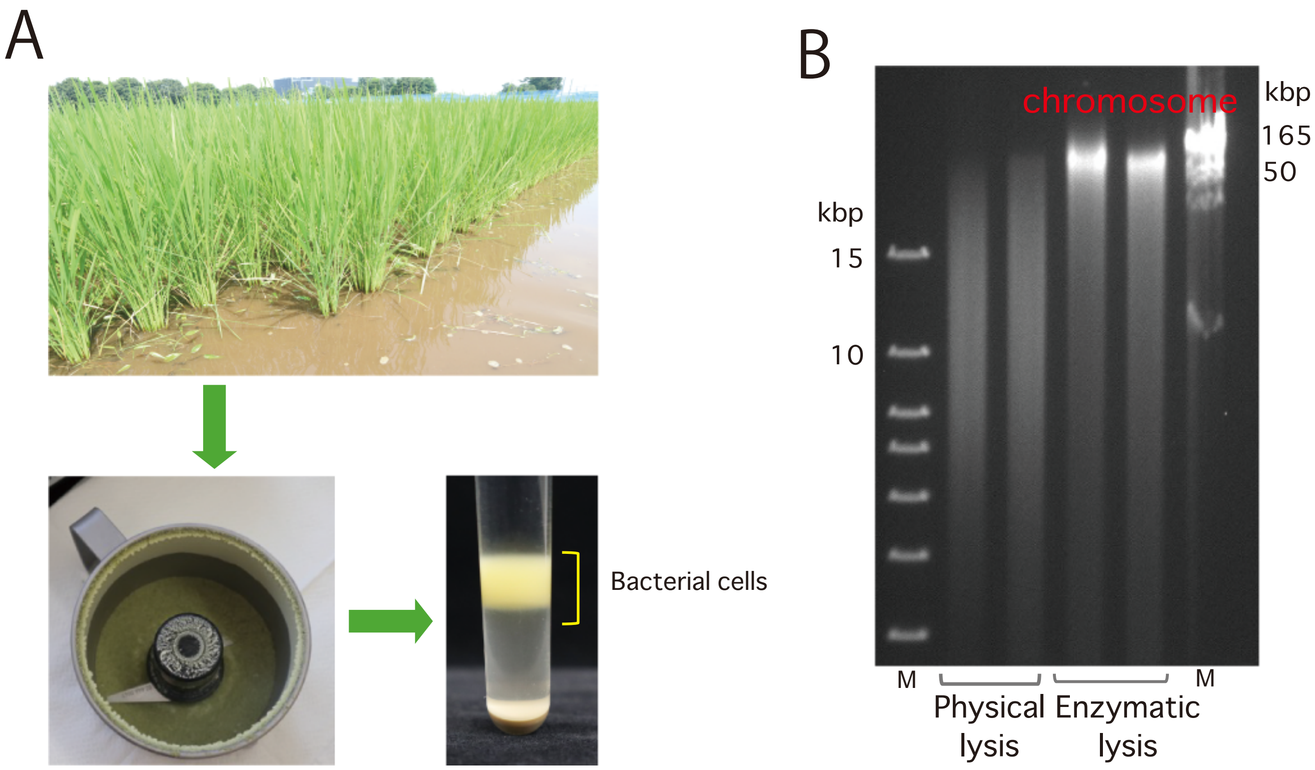


Extended Data Fig. 1

Preparation of genomic DNA from the rice-microbiome for long-read metagenomic sequencing. (A) Rice plants were sampled from experimental field and ground with dry ice. Bacterial cells were purified from aerial parts of rice plants using cell density centrifugation. (B) Genomic DNA was extracted from the purified microbiome using physical and enzymatic lysis. The presence of chromosomal DNA was confirmed using Pulse-field gel electrophoresis. M; marker

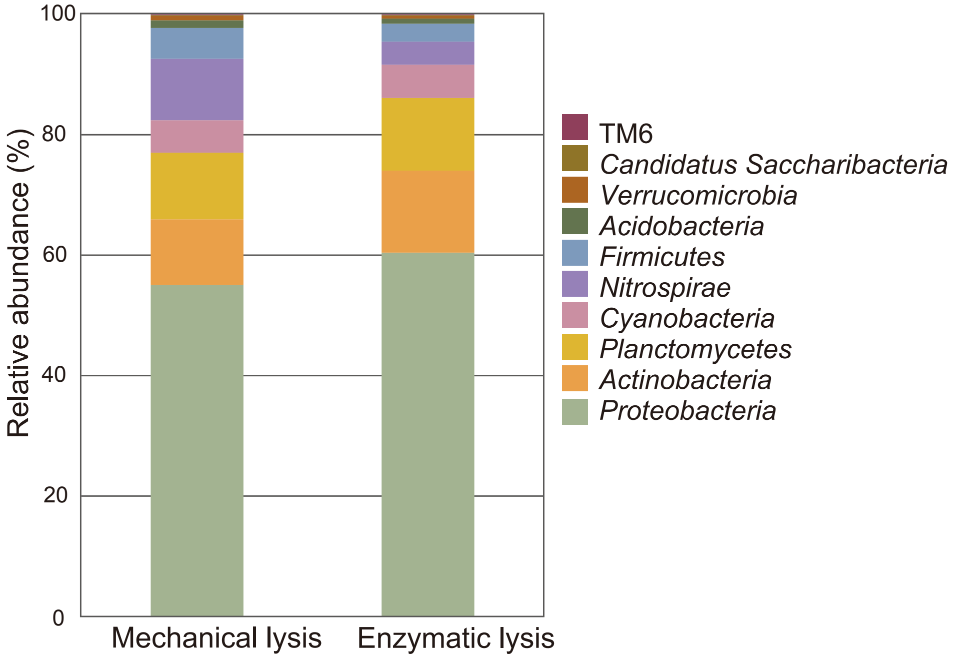


Extended Data Fig. 2

Comparison of the relative abundance of 16S rRNA genes extracted with mechanical lysis and enzymatic lysis. Each color represents the relative abundance of 16S rRNA genes at the corresponding taxonomic rank.

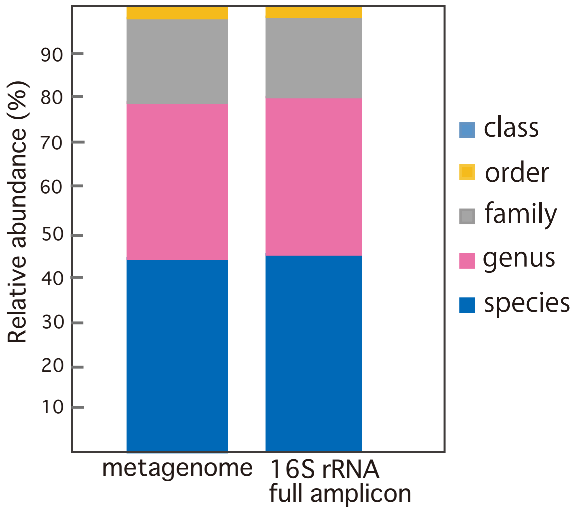


Extended Data Fig. 3

Comparison of the relative abundance of 16S rRNA genes at different taxonomical ranks in the metagenome and 16S rRNA full-length amplicon sequences. Each color represents the relative abundance of 16S rRNA genes at the corresponding taxonomic rank.

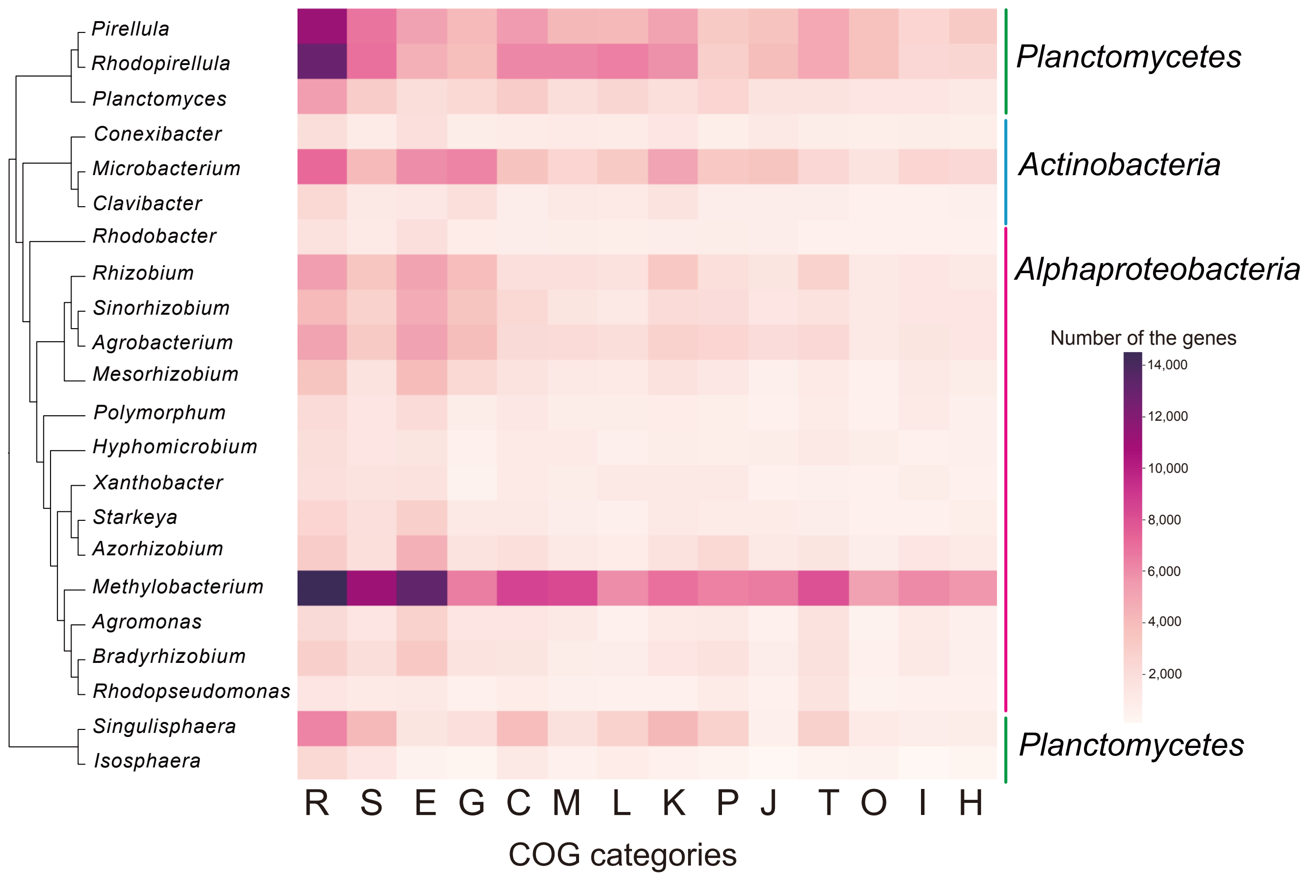


Extended Data Fig. 4

Gene categorization using the COG database. The number of genes categorized in each group is shown in the heatmap.

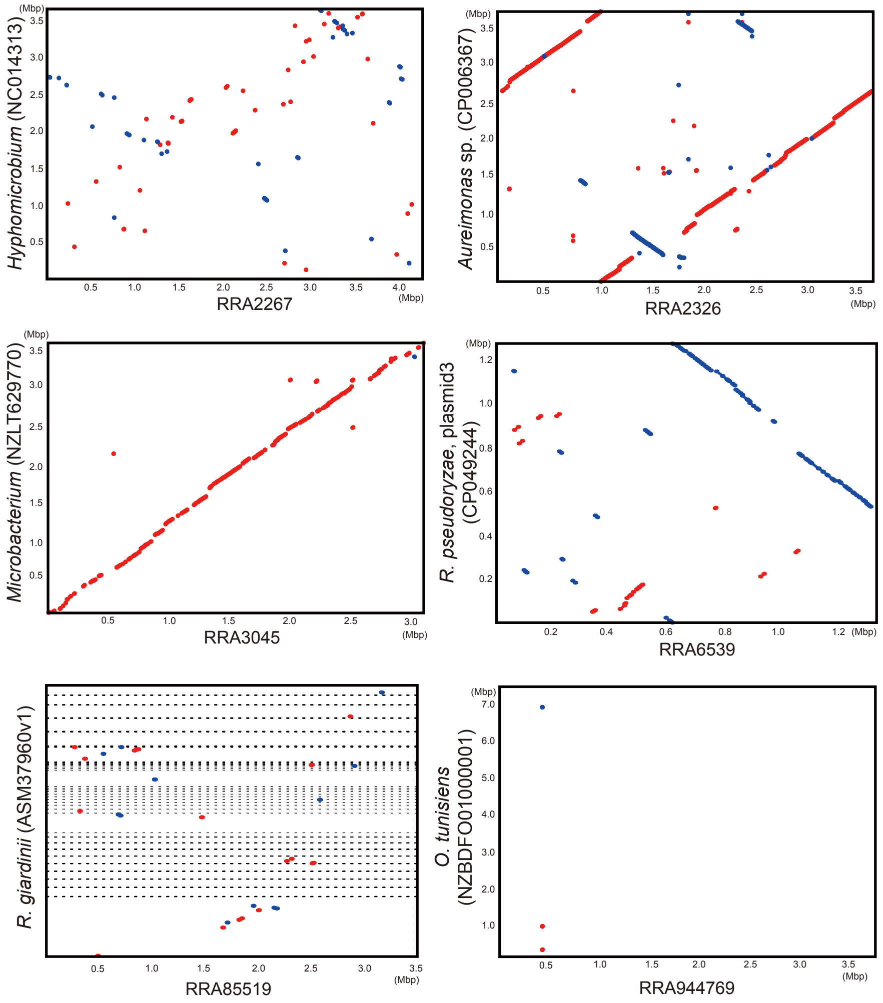


Extended Data Fig. 5

Alignment of whole genomic sequences between the six circular contigs and the closest bacterial relative genome. The genome of the reference strain, *Rhizobium glardinii*, was determined by whole genome sequencing (WGS). The bold dotted line of the horizontal axis represents each contig of *R. giardinii*.

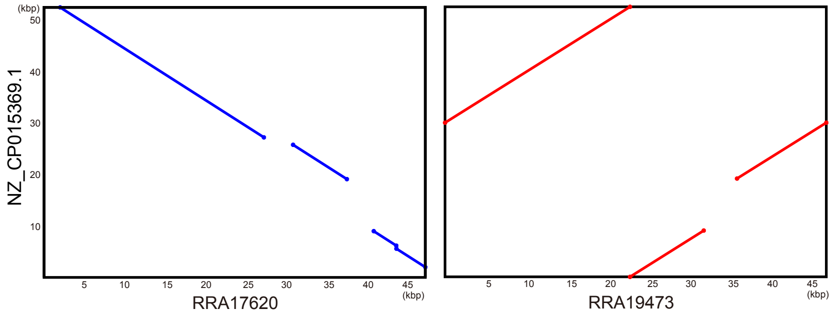


Extended Data Fig. 6

Comparison of the whole genomic sequences of RRA17620 and RRA19473 to the plasmid of *Methylobacterium phyllosphaerae* strain CBMB27 (NZ_CP015369.1).

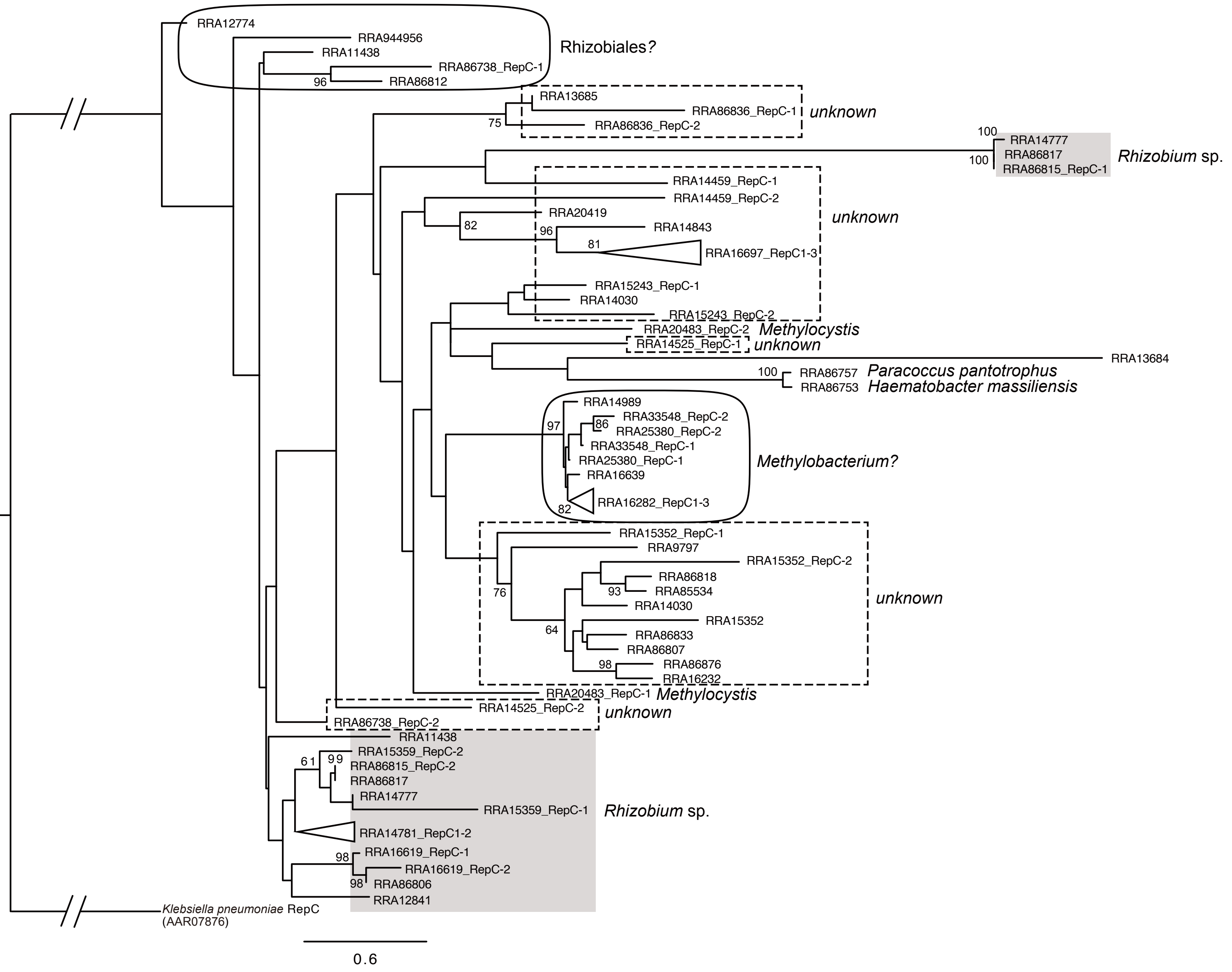


Extended Data Fig. 7

Phylogenetic tree of RepC in small circular contigs (< 1Mbp). The 61 of RepC on 39 contigs were used to construct the phylogenetic tree. The accession number indicates the representative RepC in each cluster. The taxonomy of RepC that are independently clustered with known RepC are defined as unknown. The RepC of *Klebsiella pneumoniae* (accession number: AAR07876) was used as the outgroup.

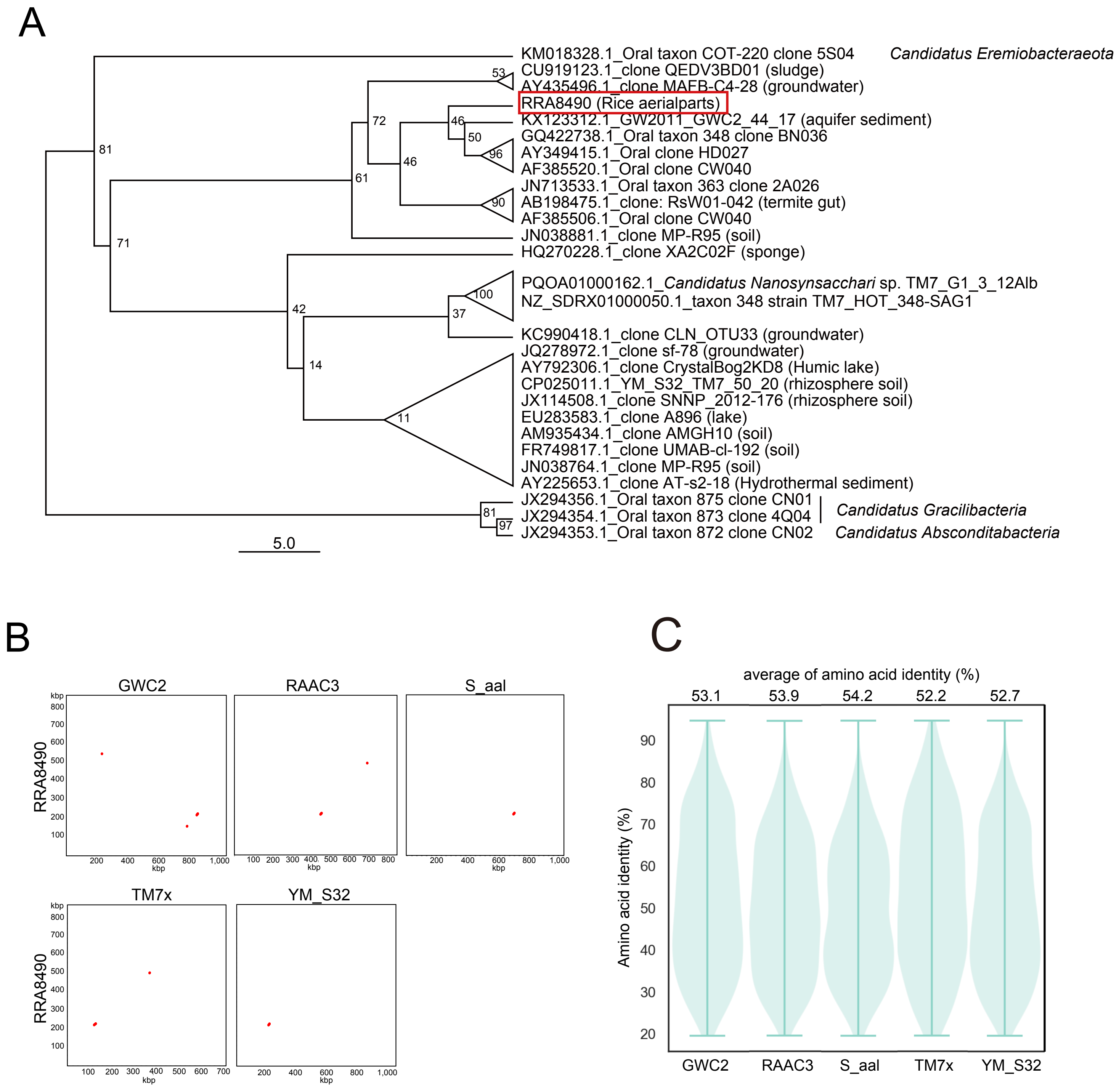


Extended Data Fig. 8

(A) a phylogenetic tree using 16S rRNA genes of strains in *Candidatus Sacchribacteria*. (B) Comparison of whole genomic sequences between RRA8490 and five strains. Whole genomic sequences were compared using nucmer, showing that RRA8490 is not similar to the others. (C) amino acid identity between RRA8490 and the five strains in *Candidatus Sacchribacteria*. The average amino acid identity was calculated using the AAI calculator in Kostas lab with default parameters (http://enve-omics.ce.gatech.edu/aai/).

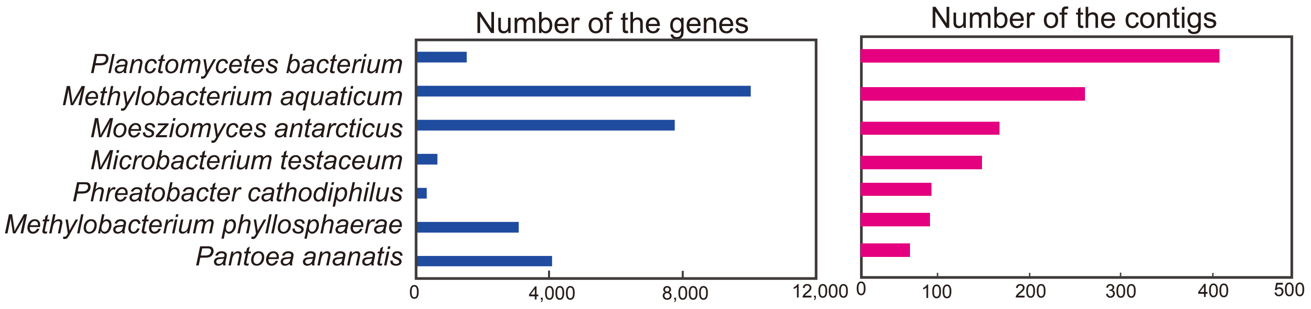


Exrended Data Fig. 9.

The number of the genes identified in metagenome with high identity and coverage to specific bacterial species and of the contigs carrying these genes. The contigs counted more than 50 were shown.
